# Evolution of shell flattening and the loss of coiling in top shells (Gastropoda: Trochidae: Fossarininae) on wave-swept rock reefs

**DOI:** 10.1101/318394

**Authors:** Luna Yamamori, Makoto Kato

## Abstract

Flattening of coiled shells has occurred in numerous gastropod lineages, probably as an adaptation to life in narrow protected spaces, such as crevices or the undersides of rocks. While several genera in the top snail family (Trochidae) have flattened shells, two Fossarininae genera, *Broderipia* and *Roya*, are unique in having shells that are limpet-like and zygomorphic, lacking any trace of coiling. The sister genera of these two genera are *Fossarina* and *Synaptocochlea*, both of which have coiled shells and live in rock crevices or the vacant shells of sessile organisms. Although *Broderipia* has recently been identified as living symbiotically in the pits of sea urchins, the habitat and biology of *Roya* are poorly known. After an extensive search for rare *Roya* snails on rocky shores of the Japanese Archipelago, we found live *Roya eximia* snails on intertidal/subtidal rock surfaces exposed to strong waves. The *Roya* snails crawled swiftly over wave-swept rock surfaces at low tide, while they retreated into the vacant shells of barnacles at high tide, where they adhered firmly to the inner wall. A survey of the macrobenthic communities around the snail habitat showed that *Roya* snails inhabited only wave-swept rocks of exposed reefs, where the substrata was covered by encrusting red algae and barnacles. Despite the abnormal shell morphology, the radula was similar to other species in the subfamily, and the diet of *Roya* snails was mainly pennate diatoms. The limpet-like shell of *Roya* caused loss of coiling and contraction of the soft body, acquisition of a zygomorphic flat body, expansion of the foot sole and loss of the operculum. All of these changes improved tolerance of strong waves and the ability to cling to rock surfaces, and thus enabled a lifestyle split between wave-swept rock surfaces and refugia of vacant barnacle shells.

## Introduction

Molluscs exhibit a wide range of shell forms as adaptations to surrounding environmental conditions [1], and for defense against predators [2]. In the history of shell-shape evolution, flattening of the coiled shell is one of the most common events; some of these flattened shells subsequently lost their coiling, forming low-conical limpet-shaped shells. Limpet-shaped shells are seen in Patellogastropoda, Cocculiniformia, Lepetodriloidae, Fissurellidae, Phenacolepadidae, Hipponicidae, Calyptraeidae, Umbraculidae, Trimusculidae, Siphonariidae, Ancylidae, some Capulidae, *Thyca crystallina* (Eulimidae), *Amathina* (Amathinidae), and a portion of Fossarininae (Trochidae), among other groups [3]. Of these taxa, only the first five lineages listed originally exhibited non-coiled shells.

Patellogastropoda is the largest taxon with limpet-shaped shells, which generally attach to rock surfaces with strong adhesive power and feed by grazing on macroalgae and benthic diatoms [4]. Cocculiniformia attach to sunken wood or whale bone in the deep sea, feeding from their adhered foundation [5-6]. Lepetodriloidae live near hydrothermal vents, with occasional grazing and active suspension feeding, and some species host filamentous bacterial episymbionts on their gills [7]. Fissurellidae snails have limpet-shaped shells with a small oval hole on top, or a small cut on the back end, of the shell. Most species of Fissurellidae are herbivorous, feeding on diatoms, cyanobacteria, macroalgae and sea grasses, whereas emarginuline and diodorine species have been reported to feed on sponges and mixed detrital materials [8-9]. Phenacolepadidae inhabit the bottom surfaces of deeply-embedded rocks or decaying wood, which can be described as a dysoxic, sulfate-rich environment. Phenacolepadid gastropods are thought to feed on chemosynthetic bacteria [10]. Hipponicidae attach to hard inorganic or organic substrata such as rocks, dead corals, and the shells of large gastropods by extracting calcareous substances, and are filter feeders [11]. Calyptraeidae normally attach to solid organic substrates such as dead bivalves, shells occupied by hermit crabs, and the undersides of horseshoe crabs. Calyptraeids are filter feeders that consume particulate foods in water, including small plankton, detritus, and excrement of their host [12]. Umbraculidae have vestigial flat shells that are deeply coated with mantle, feeding on sponges [13]. Trimusculidae gather particles of phytoplankton using a mucus curtain secreted from glands on the head, taking advantage of the turbulence of water [14]. Siphonariidae is an air-breathing family with asymmetrical limpet-shaped shells, which feed on microalgae such as cyanobacteria and diatoms [15]. Ancylidae are limpet-shaped freshwater snails that attach to rocks or aquatic plants, feeding on diatoms on their substrates [16]. The limpet-shaped Capulid genus *Capulus,* which have top-curled limpet-like shells, attach mainly to bivalves and exploit the feeding currents of their host bivalves or steal the bivalves’ accumulated food using a pseudoproboscis [17-18]. *Thyca crystallina* is an obligate parasite of sea star *Linckia* spp., feeding on its host’s hemal and perihemal fluids [19]. *Amathina* snails with top-curled limpet-like shells attach to large bivalves, such as the fan shell *Pinna bicolor*, and collect waste food materials from their host bivalves by inserting their proboscis inside the host’s shell [20].

Branch (1985) presented some major advantages of the limpet-shaped shell [3]. First, conical shells greatly reduce water resistance, such that limpets can venture into strongly wave-exposed areas where most coiled gastropods cannot maintain adherence. Second, the large aperture of the limpet-shaped shell allows development of a large foot. Limpets cannot withdraw into their shells or protect themselves with an operculum, instead utilizing a strong clinging force with hard substrata, which is derived from the large foot, as protection from predation. However, the factors that promote the evolution of limpet-like shells have not yet been identified, because most members of the aforementioned families have limpet-like shells and species in transition from coiled to non-coiled shells are rare.

The top-shelled family Trochidae is characterized by conical, coiled shells and an alga-grazing habit, although some linages are filter feeders (Umboniinae) and others have flattened shells (Alcyninae, Fossarininae and Stomatellinae). In particular, a completely limpet-like shell is seen only in two genera of Fossarininae, *Broderipia* and *Roya.* The sister genera of *Broderipia* and *Roya* are *Fossarina,* with a round-spiral shell and *Synaptocochlea* with a loosely coiled auriform shell like abalone [21]. Because the four genera of Fossarininae are currently at various stages in the evolutionary process of shell flattening, comparisons of their biology and habitats may help clarify the selection pressures driving shell flattening and loss of coiling. *Broderipia*, which has an extremely flat shell, was recently revealed to be symbiotic in the pits of sea urchins, and its flat limpet-like shell is apparently adaptive to life in the narrow open space of the pits [22]. On the other hand, the habitat and biology of *Roya* are poorly understood.

To detect selective pressures acting on shell flattening and the loss of coiling in Fossarininae, we first conducted an extensive search for habitats of the key genus, *Roya*. Because *Roya* snails are found in low intertidal areas of wave-swept rocky reefs, we carried out a field survey of the macrobenthic and macrophytic communities on various types of rocky reefs surrounding *Roya* snails, and observed the diurnal behavior of the snails. To determine their feeding biology, radula and gut contents of *Roya* snails were also examined. By superimposing the data thus obtained on a phylogenetic tree of Fossarininae, we clarified the evolution of shell flattening and loss of coiling in gastropods.

## Materials and Methods

### Study sites

This study was conducted on rocky shores in the Japanese Archipelago, which are influenced by the warm Kuroshio Current. Through a preliminary search for the rare limpet-like trochid snail *R. eximia,* five populations were identified on the southern coasts of the Kii Peninsula and Shikoku Island (sites A–E in Fig 1). At all sites, the maximum tidal range during spring tide is around 200 cm. Because the habitat of *R. eximia* is constantly exposed to violent waves, accessible sites are rare. The only accessible coastal sites were sites A (33°69′51′′N, 135°33′58′′E) and E (32°46'00"N, 132°37'18"E). Site A is a small near-shore island called Toshima, the west coast of which faces the Kii Channel and is exposed to strong waves. The rock bed is conglomerate mixed with brittle sandstone, and the surface remains rough due to constant erosion from waves (Fig 2a–b). Site E is a rocky and boulder-filled shore of Kashiwajima Island, which is constantly exposed to strong waves. The large boulders accumulated on the shore are made up of hard igneous rock [23](National Institute of Advanced Industrial Science and Technology 2018), the surface of which is smooth (Fig 2c–d).

**Fig 1.**
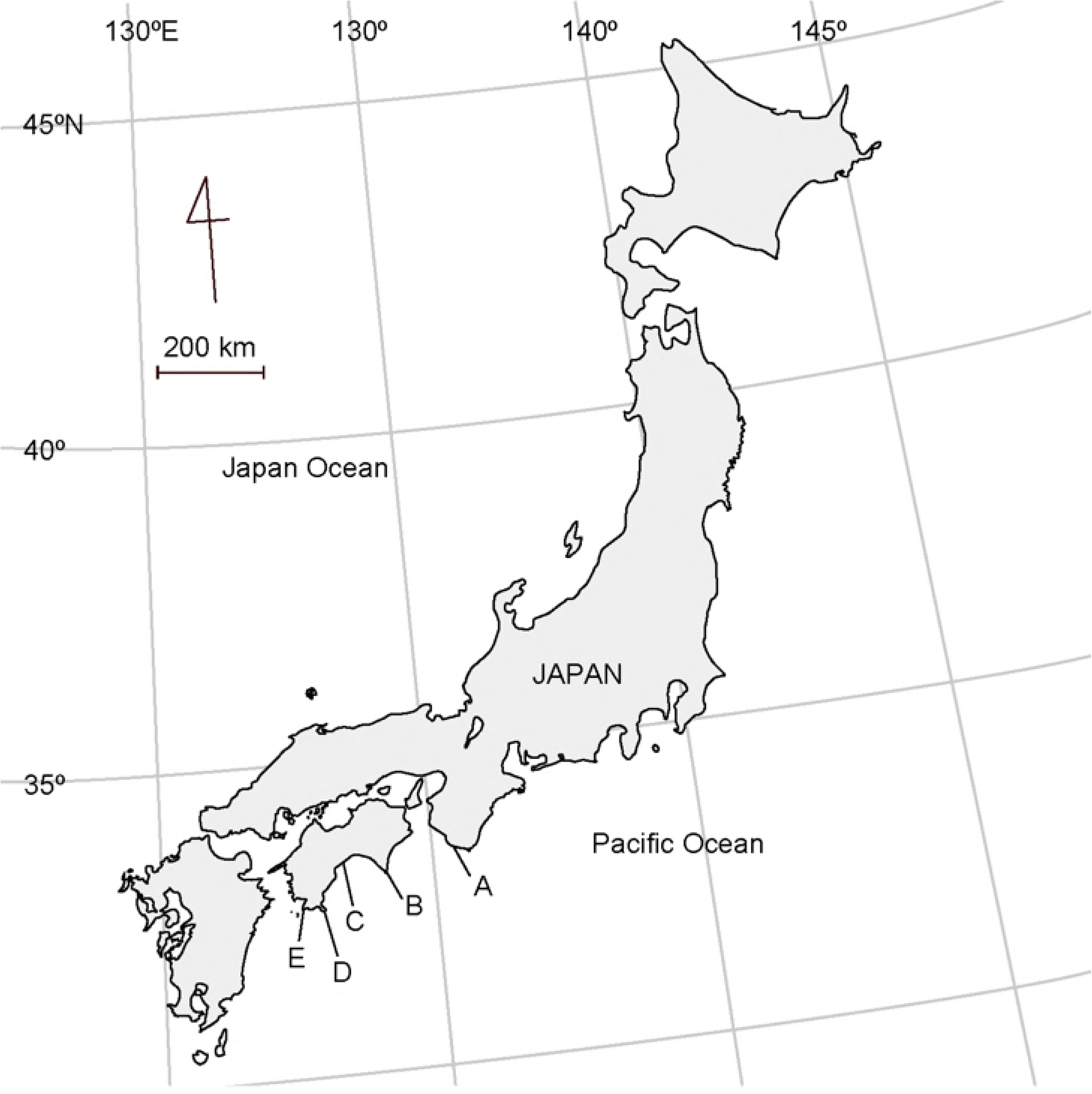
Locations of the study sites. A: Shirahama in Wakayama Prefecture, B: Muroto Cape, C: Goshikihama, D: Chihiro Cape, E: Kashiwajima Island in Kochi Prefecture.

**Fig 2.**
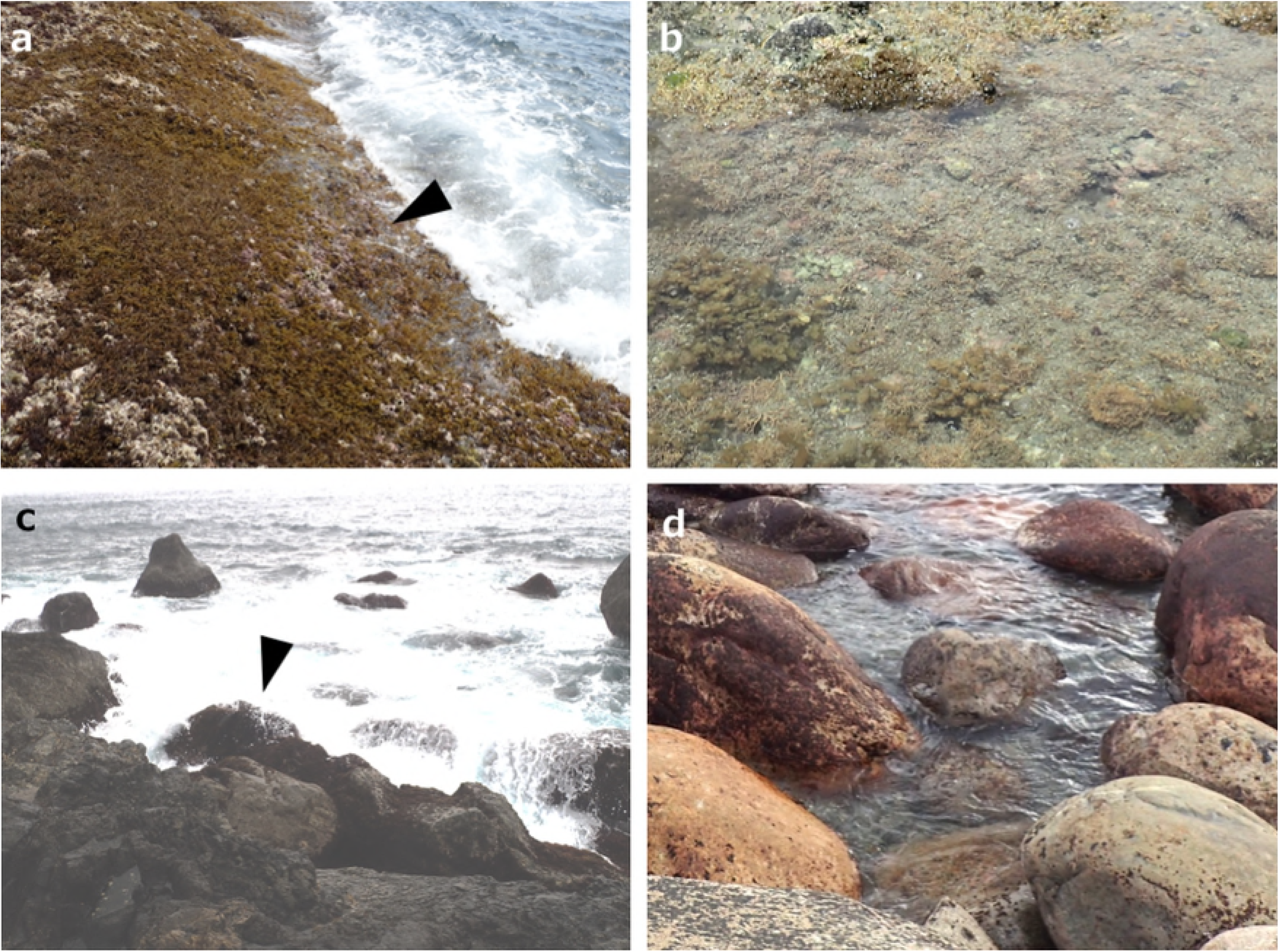
Study sites and habitats of snail species belonging to Fossarininae. a: Exposed reef at site A; arrowhead shows the tidal level inhabited by *R. eximia*. b: Protected reef at site A. c: Exposed reef at site E; arrowhead shows the rock inhabited by *R. eximia*. d: Protected reef at site E.

### Census on macroalgae and macrobenthos

In the lower intertidal zones of sites A and E, two different settings of rock surfaces were chosen, which were exposed to and protected from strong waves. We refer to the former and latter settings as exposed and protected reefs, respectively. In both rock surface settings, 10 quadrats (10 × 10 cm) were set, and all macroalgae and macrobenthos in each quadrat were examined in terms of species, abundance (for macrobenthos) and coverage (for macroalgae) during a spring tide in April 2017.

### Morphology and diet of Fossarininae

To compare the shell and radula morphologies of the four Fossarininae species, the side view of the living snail and side view and aperture view of their shells were photographed. Radulae were removed from each species and examined using an electron microscope. To determine the diet of Fossarininae snails, the stomach and intestinal contents were examined. Fossarininae snails were collected at site A and immediately fixed in 4% formalin solution. After 12 h of fixation, snails were removed from the formalin solution and rinsed with flowing fresh water for 30 minutes. Then, the stomach and intestine of each Fossarininae snail was removed and opened in water; approximately 1/5 of the contents were observed and its organic matter was counted using an optical microscope. The rest of the material was dried in an incubator at 50 °C, and then placed on conductive carbon tape and examined carefully using an electron microscope.

### Diurnal behavior of *R. eximia*

At site A, diurnal changes in the distribution of *R. eximia* snails were surveyed in the intertidal area at high tide, the middle of the ebb tide (awash time of their habitat tidal level) and low tide in the daytime, and at low tide in the nighttime on 15^th^ April 2017. In each census, *R. eximia* snails were sought out not only on the rock surface, but also in the interspaces of sessile organisms and inside vacant shells of barnacles; their behavior was observed.

## Results

### Rock-surface environment and macrobenthic community at each site

The surfaces of the exposed reefs in the lower intertidal zones of sites A and E were covered with 13 species of encrusting and branching algae. The proportions of bare surface, i.e., areas without cover of macroalgae or sessile organisms, of exposed and protected rock surfaces were 8.86 % and 47.1 % at site A, and 24.3 % and 20.8 % at site E, respectively. Most rock surfaces were not covered with sand, although the protected rock surface of site A was partly covered with sand.

The most frequent and dominant macroalgae were the encrusting coralline red alga *Lithophyllum* spp. and encrusting non-coralline red alga *Hildenbrandia rubra* (Table 1). The branched coralline algae *Amphiroa beauvoisii* and *Serraticardia maxima* were found only at site A. The assemblages of brown algae differed between the two sites, as well as between exposed and protected microhabitats. At site A, the rock surface exposed to strong waves was dominated by the large brown alga *Sargassum fusiforme* and *S. patens*, while more protected rock surfaces were often inhabited by the brown alga *Palisada intermedia*. On the other hand, at site E, exposed rock surfaces were inhabited by *P. intermedia* and *Ishige okamurae*, while no brown algae were observed on protected rock surfaces.

The coverages of macroalgae and sessile organisms at sites A and E on exposed and protected rock surfaces are shown in Fig 3. The coverage of branching red algae on exposed rock surfaces was greater at site A than site E (t-test, p=0.011), whereas the coverage of encrusting red algae on protected rock surfaces was greater at site E than site A (t-test, p=0.0047). Coverages of encrusting red algae and branching brown algae at site A were significantly greater on exposed rock surfaces than on protected rock surfaces (t-test: p=0.033 for encrusting red algae; p=0.047 for branching brown algae). In addition to these macroalgae, the rock surfaces of the intertidal zones were covered with sessile organisms, such as barnacles and a sessile vermetid snail, *Serpulorbis imbricatus*. At site A, three large barnacle species, *Megabalanus volcano*, *Tetraclita japonica* and *T. squamosal* were observed on exposed rock surfaces, while no barnacles were found on protected rock surfaces. At site E, four barnacle species were observed. Two species of small barnacles, *Fistulobalanus albicostatus* and *Chthamalus challengeri,* inhabited the protected rock surfaces, while three species, including the two large tetraclitid species *T. japonica* and *T. squamosal,* inhabited exposed rock surfaces (Table 1). Coverage of sessile organisms on exposed rock surfaces was greater at site E than site A (t-test, p=0.00050); at site E, coverage was greater on exposed rock surfaces than protected rock surfaces (t-test, p=0.00032).

**Fig 3.**
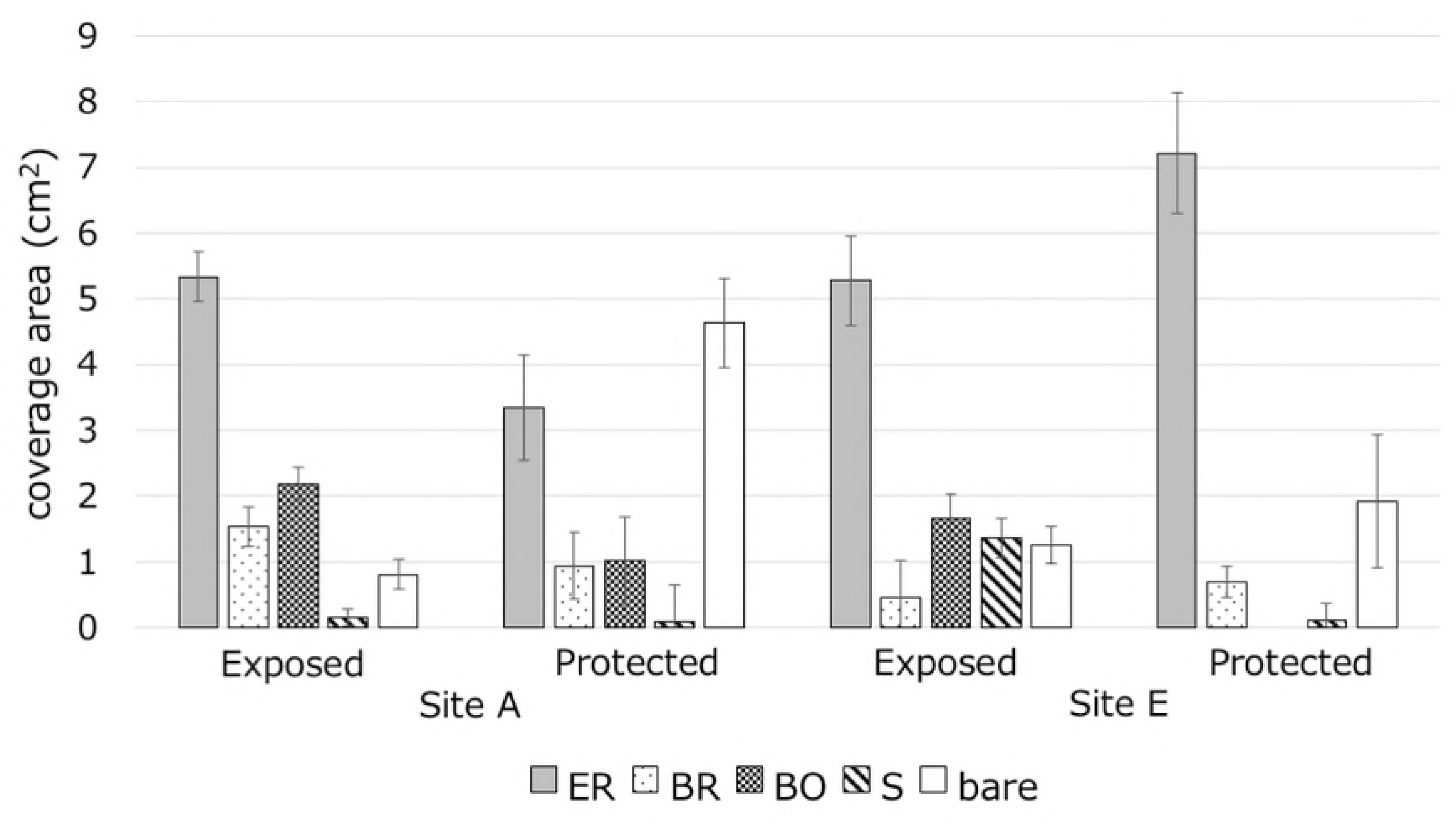
Mean coverage (cm^2^) of various algae and sessile organisms in 10 × 10 cm quadrats (n = 10) set on exposed and protected rock surfaces at sites A and E. ER: encrusting Rhodophyta, BR: branching Rhodophyta BO: branching Ochrophyta. S: sessile invertebrates, bare: bare surface

**Table 1.**
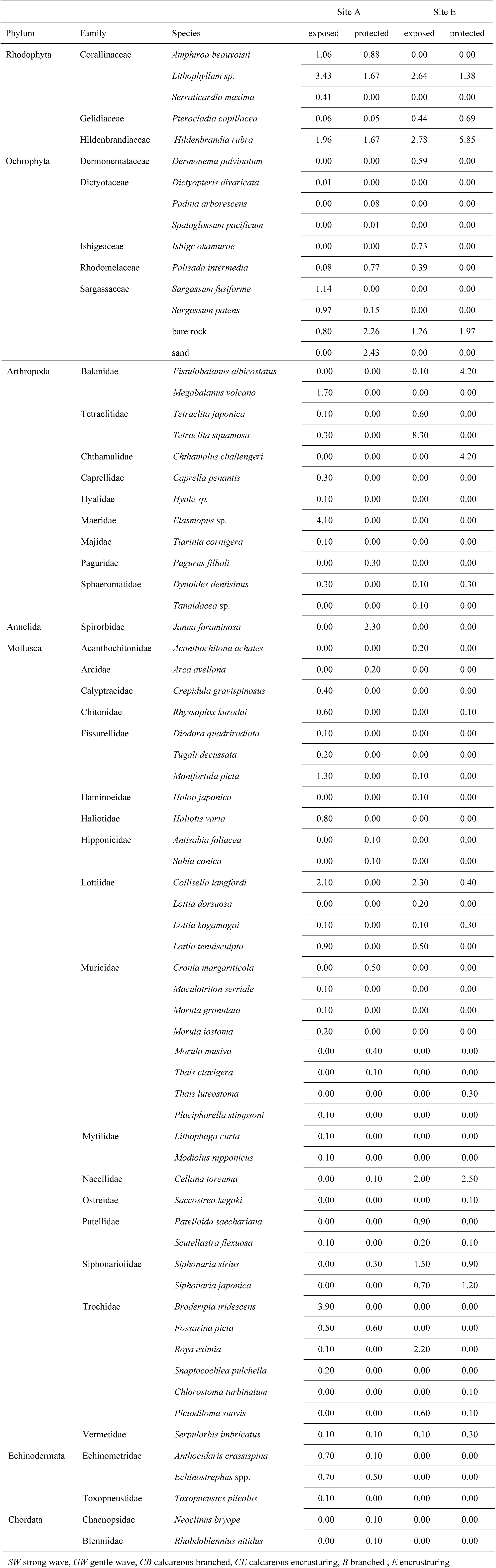
Species found in quadrats and degree of coverage of each macroalgal species (n = 10)

Both the species richness and density were compared between exposed and protected reefs (Fig 4). Species richness on exposed habitats was higher at site A than at site E (t-test, p=0.018), and species richness on the exposed reef was higher than on the protected reef at both sites A and E (t-test: p=0.000011 for site A, p=0.00040 for site E). The density of macrobenthos on the exposed reef was also greater than that on the protected reef (t-test: p=0.024 for site A, p=0.006 for site E).

**Fig 4.**
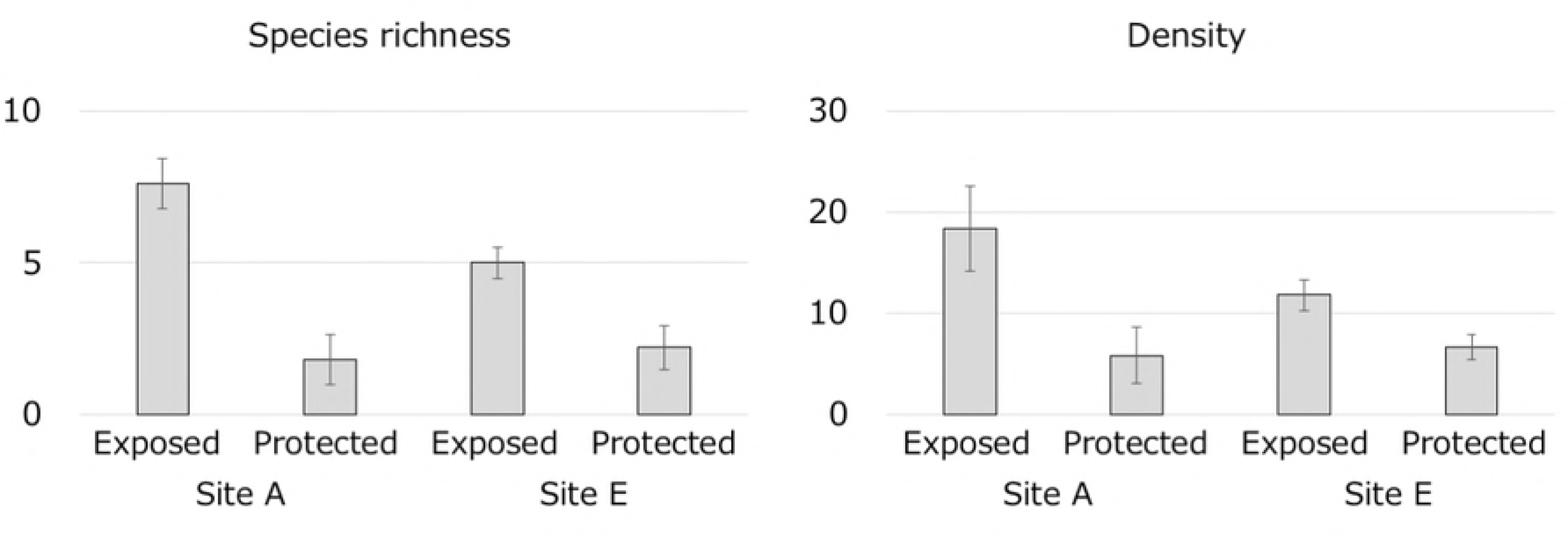
Species richness and density of the macrobenthos in 10 × 10 cm quadrats (n = 10) set on exposed and protected rock surfaces at sites A and E.

At site A, four Fossarininae species were observed in four microhabitats: open rock surfaces, crevices, sea urchin pits and vacant barnacle shells (Table 2). *Fossarina picta* was found on both protected and exposed reefs, specifically on open rock surfaces of the protected reef (Fig 5a) and inside crevices and vacant shells of barnacles on the exposed reef (Fig 5b–c). *Synaptocochlea pulchella* was found in crevices on rocks in the exposed reef (Fig 5d–e). *Broderipia iridescens* was observed exclusively inside the pits or crevices inhabited by sea urchins (Fig 5f–g). *R. eximia* was found on bare rock surfaces around barnacle colonies in exposed area (Fig 5h). The density of *R. eximia* at site E was significantly greater than at site A (t-test, p=0.033).

**Fig 5.**
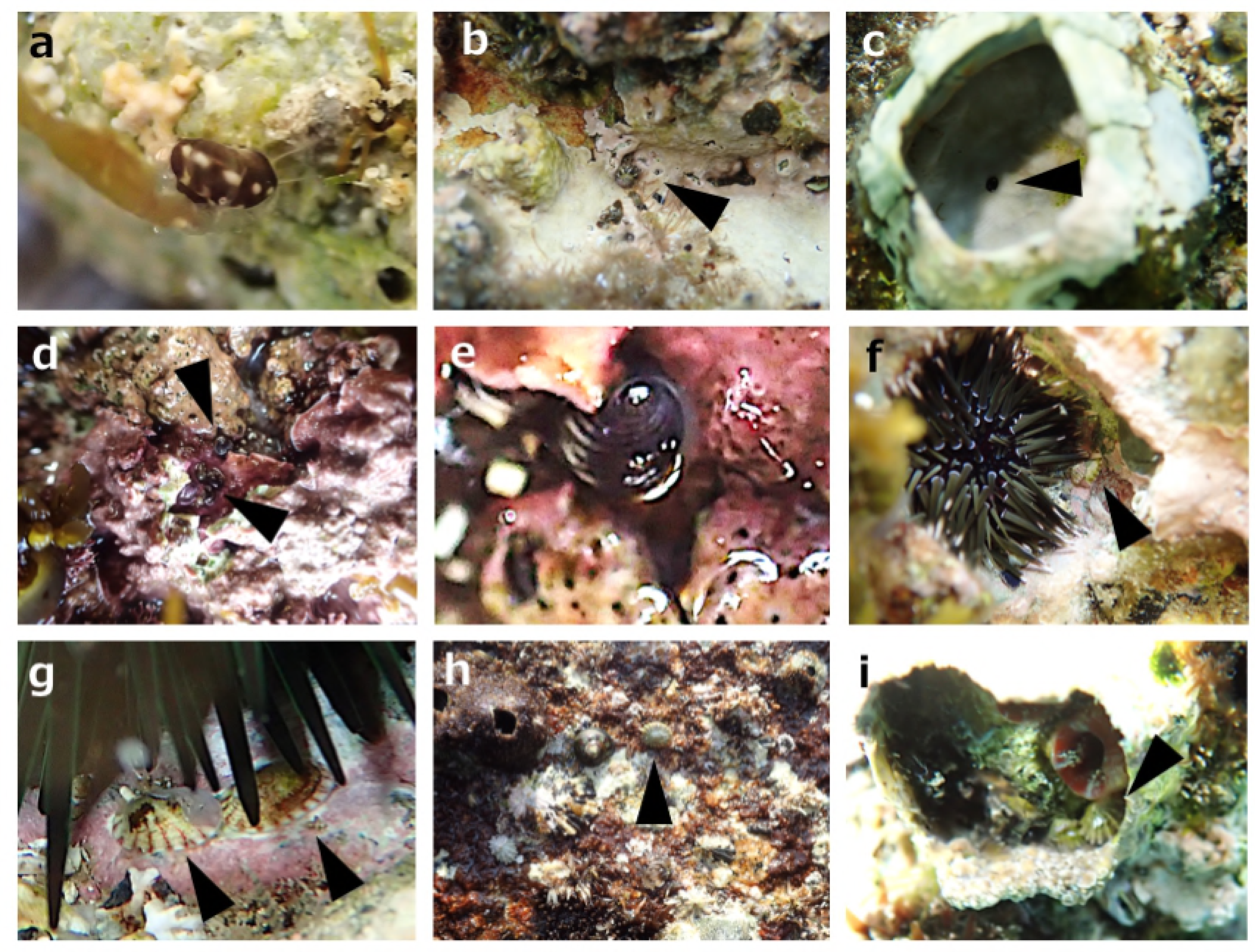
Microhabitats of Fossarina snails. a–c: *F. picta* in rock crevices of both protected and exposed rock reefs (a, b); sandy sediments are sometimes accumulated in crevices, as shown in a. *F. picta* also live in vacant shells of the barnacle *Megabalanus volcano* on exposed reefs (c). d–e: *S. pulchella* in a rock crevice. f–g: *B. iridescens* in sea urchin pits. h–i: *R. eximia* found on wave-swept rock surfaces at low tide (h), and in vacant shells of the barnacle *M. volcano* at high tide (i).

**Table 2.** The numbers of individuals of the four Fossarininae species observed at low tide.

|  | Open rock surface | Crevice | Sea urchin's pit | Vacant barnacle shell | Total |
| --- | --- | --- | --- | --- | --- |
| *F. picta* | 6 | 2 | 0 | 3 | 11 |
| *S. pulchella* | 0 | 2 | 0 | 0 | 2 |
| *B. iridescens* | 0 | 0 | 39 | 0 | 39 |
| *R. eximia* | 23 | 0 | 0 | 0 | 23 |

### Diurnal behavior of *R. eximia*

Diurnal changes in the number of *R. eximia* snails found at site A are shown in Fig 6. The habitat of *R. eximia* was within 50 cm of the ebb tide line. This habitat emerges from the water for about 1 hour during the spring tide, and is constantly splashed during its emergence. During the daytime high tide, four *R. eximia* were found inside vacant shells of the barnacle *M. volcano*, completely covered with water (Fig 5i), and all four snails had tightly adhered their bodies to the inner surface of the barnacle shells, and did not move. At mid-ebb tide in the daytime, on the other hand, the snails swiftly crept over rock surfaces awash with seawater and grazed. When tide was low during the daytime, six snails were observed on the wet surface of emergent rocks. Two snails slowly and intermittently moved over the emergent zone and grazed, while the other four snails remained motionless, with their bodies tightly adhered to the rock surface. At the nighttime low tide, three snails were observed remaining motionless on the wet rock surface.

**Fig 6.**
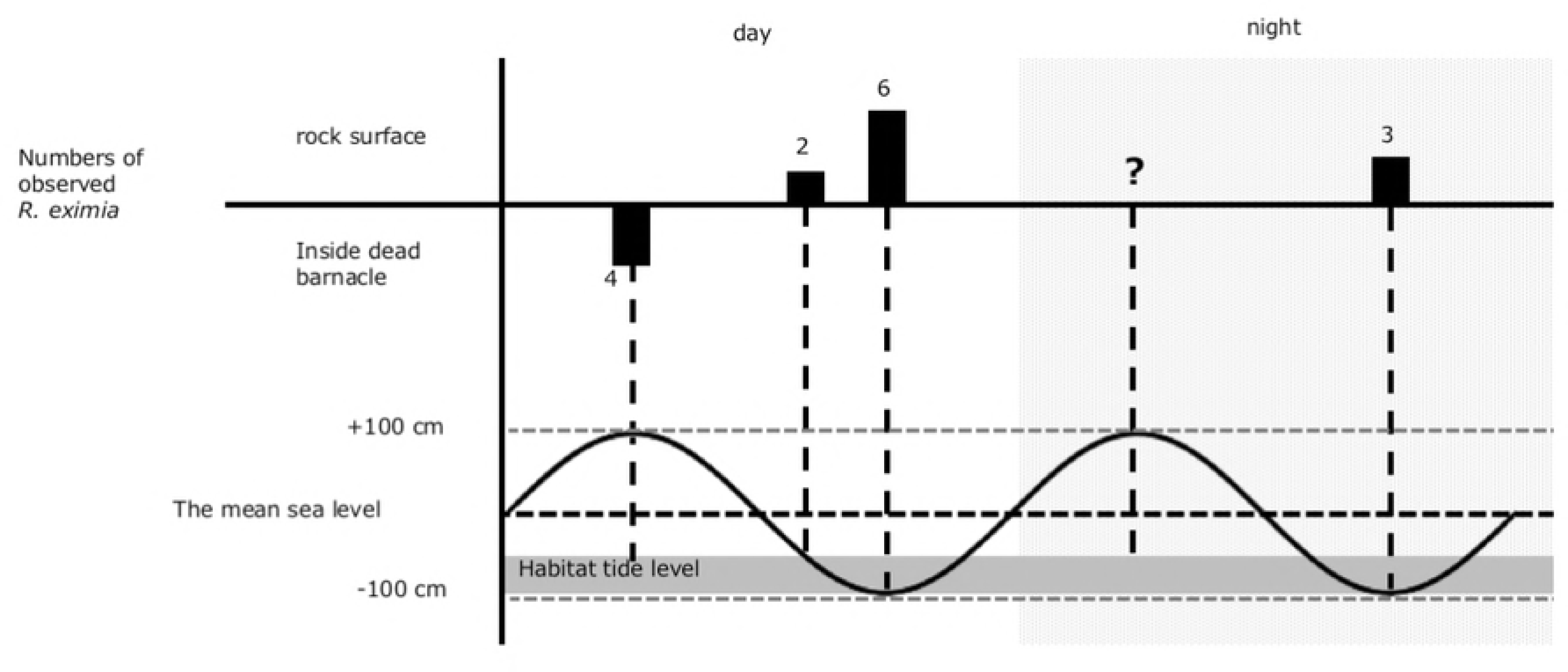
Diurnal changes of tidal level (wavy line) and observed number of *R. eximia* snails (solid columns). The habitat of the snail in reference to the tidal level is illustrated as a gray bar. *Roya* snails were observed on wet rock surfaces at low tide, while they were found inside vacant barnacle shells on submerged rock surfaces at high tide.

### Morphology of the shell, soft body and radula

The side view of a living snail, and side and aperture views of shells, are shown in Fig 7. *F. picta* have round spiral shells and four pairs of epipodial tentacles (Fig 7 a–c). *S. pulchella* have slightly flattened shells and four pairs of epipodial tentacles (Fig 7 d–f). *B. iridescens* have completely limpet-like shells and three pairs of epipodial tentacles (Fig 7 g–i). *R. eximia* (Fig 7 j–l) also have completely limpet-shaped shells and three pairs of short epipodial tentacles. *R. eximia* have a radial rib on the shell. The first and second epipodial tentacles of *B. iridescence* are longer than those of *R. eximia*. The lengths of the epipodial tentacles from the anterior end were 2.7, 2.2, and 1.7 mm for *B. iridescens* (8.2 mm shell length), and 1.7, 1.7, and 1.6 mm for *R. eximia* (9.5 mm shell length). The eye size of *R. eximia* is smaller relative to its shell length than in the other three Fossarininae species (Fig 7a, d, g, j).

**Fig 7.**
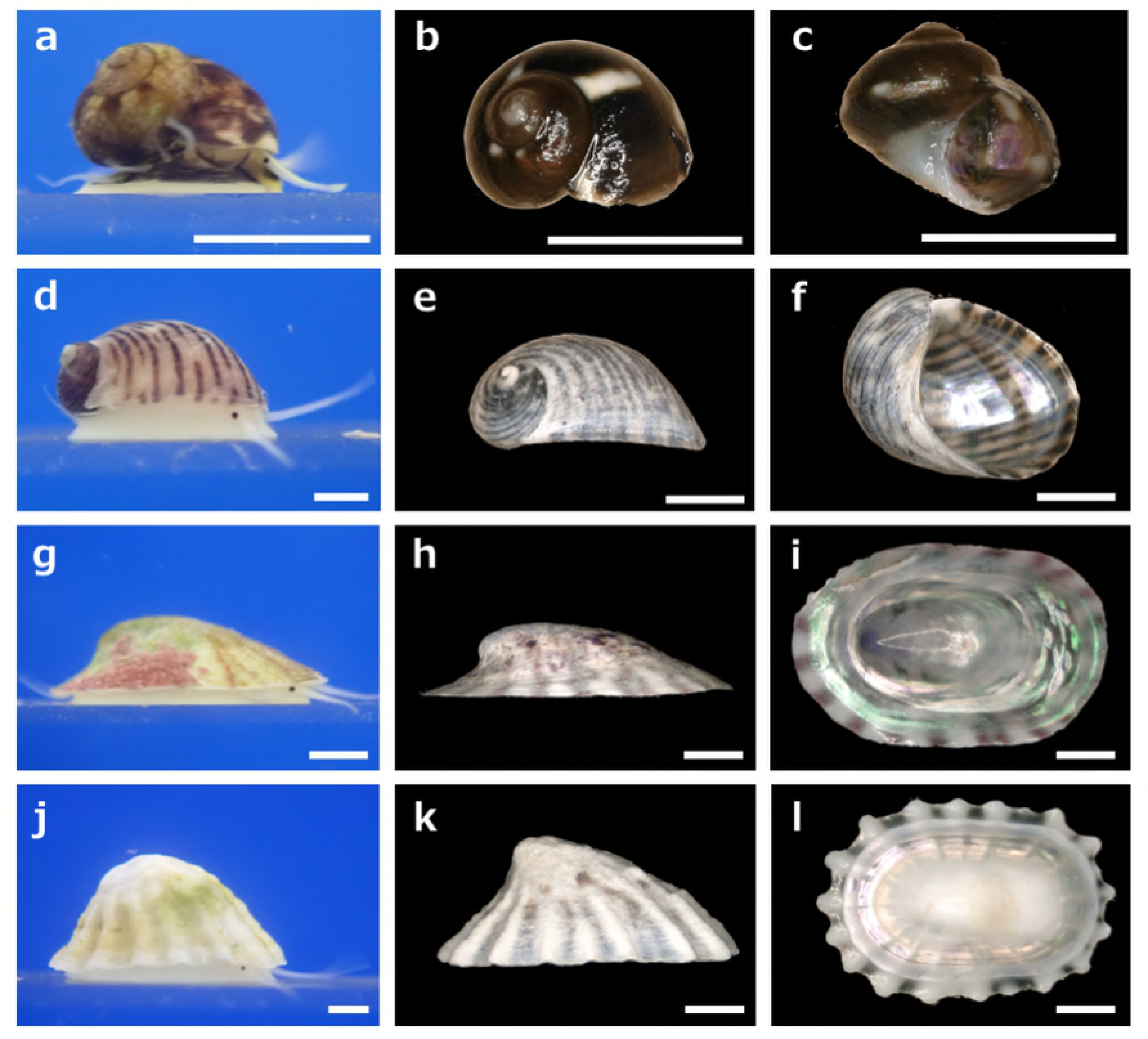
Side views of living snails (left column), and side and ventral views (middle and right columns) of shells of four Fossarininae species. *F picta* (a–c); *S. pulchella* (d–f); *B. iridescens* (g–i); *R. eximia* (j–l). *Scale bar* = 1 mm.

While the soft body of *F. picta* was coiled, the coiling was looser in *S. pulchella*, while the soft bodies of *B. iridescens* and *R. eximia* were contracted and had completely lost coiling (Fig 8). In the non-coiled soft bodies of *B. iridescens* and *R. eximia*, well-developed shell muscles are seen, and gills were only located on the left side, as in *Fossarina* and *Synaptocochlea*. Only *F. picta* and *S. pulchella* possessed an operculum. While the operculum fitted the aperture in *F. picta* (Fig 7e), the operculum of *S. pulchella* was far smaller than the large aperture of the snail (Fig 7f). The radulae of all four Fossarininae species are rhipidoglossate. The central and lateral teeth of *F. picta* clearly differ from those of the other three species, whereas the marginal teeth were very similar in all of four species. The formula of *F. picta* radula is (30–40)–4–1–4–(30–40) (Fig 9a). The central tooth has no triangular edge and lies flat along the radula, with the cusp curled slightly upward. The teeth become much narrower in the basal region (Fig 9b). The lateral teeth on each side are hidden by the central tooth or the adjacent lateral tooth. The size and shape of the lateral teeth are similar to those of the central tooth, but the lateral teeth are slightly asymmetric with the outer side being wider than the inner side. The marginal teeth are oar-shaped, with 4–6 serrations on each side (Fig 9c). The formula of *S. pulchella* radula is (30–40)–5–1–5–(30–40) (Fig 9d). The central tooth has a sharp triangular edge with 2–3 serrations on each side (Fig 9e). The lateral teeth on each side are similar in size and shape to the central teeth, and bent toward the center of the radula. The marginal teeth are oar-shaped with serrations (Fig 9f). The formula of *B. iridescens* radula is (30–40)–5–1–5–(30–40) (Fig 9g). The central tooth has a shovel-like triangular edge with 4–6 serrations on each side (Fig 9h). The lateral teeth on each side are similar in size and shape to the central teeth, and are bent toward the center of the radula. The marginal teeth are oar-shaped and serrations are present on all teeth (Fig 9i). The formula of *R. eximia* radula is (30–40)–7–1–7–(30–40) (Fig 9j). The shapes and sizes of all teeth of *R. eximia* are very similar to those of *B. iridescens*, but the cutting edges of the teeth are slightly sharper (Fig 9k–l).

**Fig 8.**
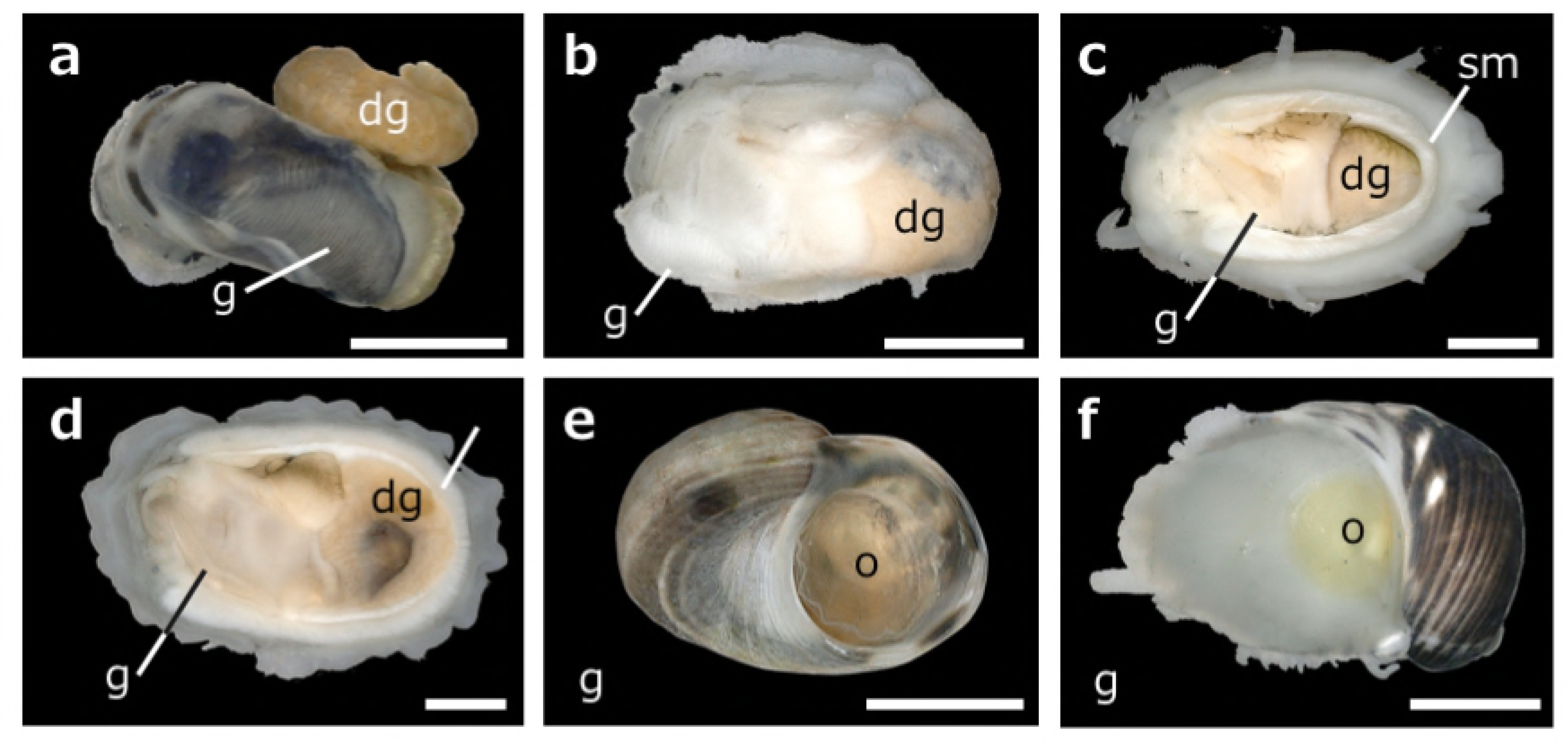
Dorsal views of the soft bodies (a–d) and opercula (e–f) of four Fossarininae species. *F. picta* (a, e); *S. pulchella* (b, f); *B. iridescens* (c); *R. eximia* (d). *g* gill, *dg* digestive gland, *sm* shell muscle, *o* operculum. *Scale bar* = 1 mm.

**Fig 9.**
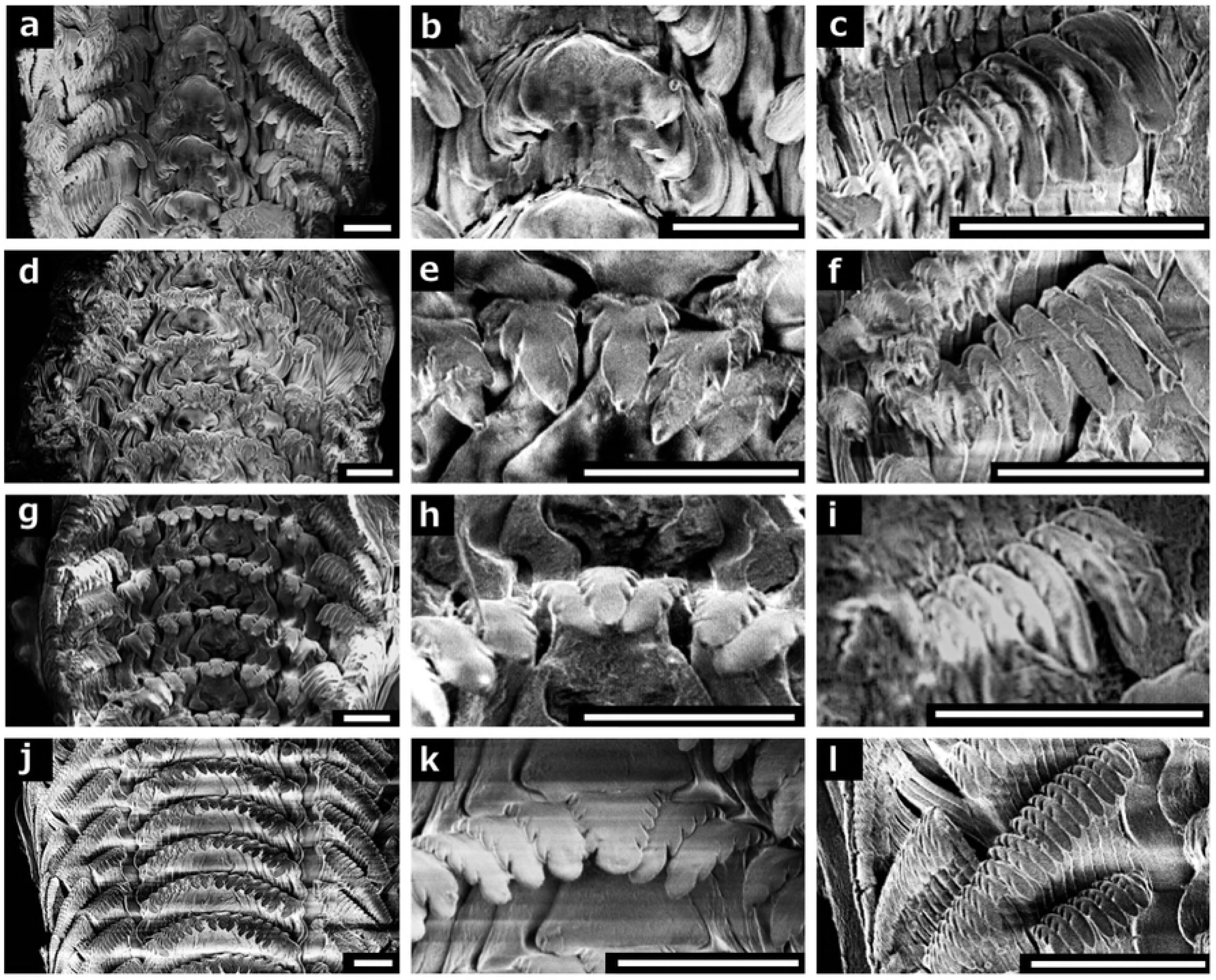
Views of the whole radulae (left column) and close-up views of the central and marginal teeth (middle and right columns) of four Fossarininae species. *F. picta* (a–c); *S. pulchella* (d–f); *B. iridescens* (g–i); *R. eximia* (j–l). *Scale bar* = 30 μm.

### Diet of Fossarininae

Diatoms were the primary component of the gastric contents of all four Fossarininae species (Fig 10a, c–f; Table 3). Fragments of red algae (Fig 10b) were also observed in small amounts in all snail species, while nematodes and sponge ossicles were rarely observed. Diatoms in order Pennales, shown in Fig 10c, represented a large proportion of the gastric contents in all four snail species, especially *F. picta*.

**Fig 10.**
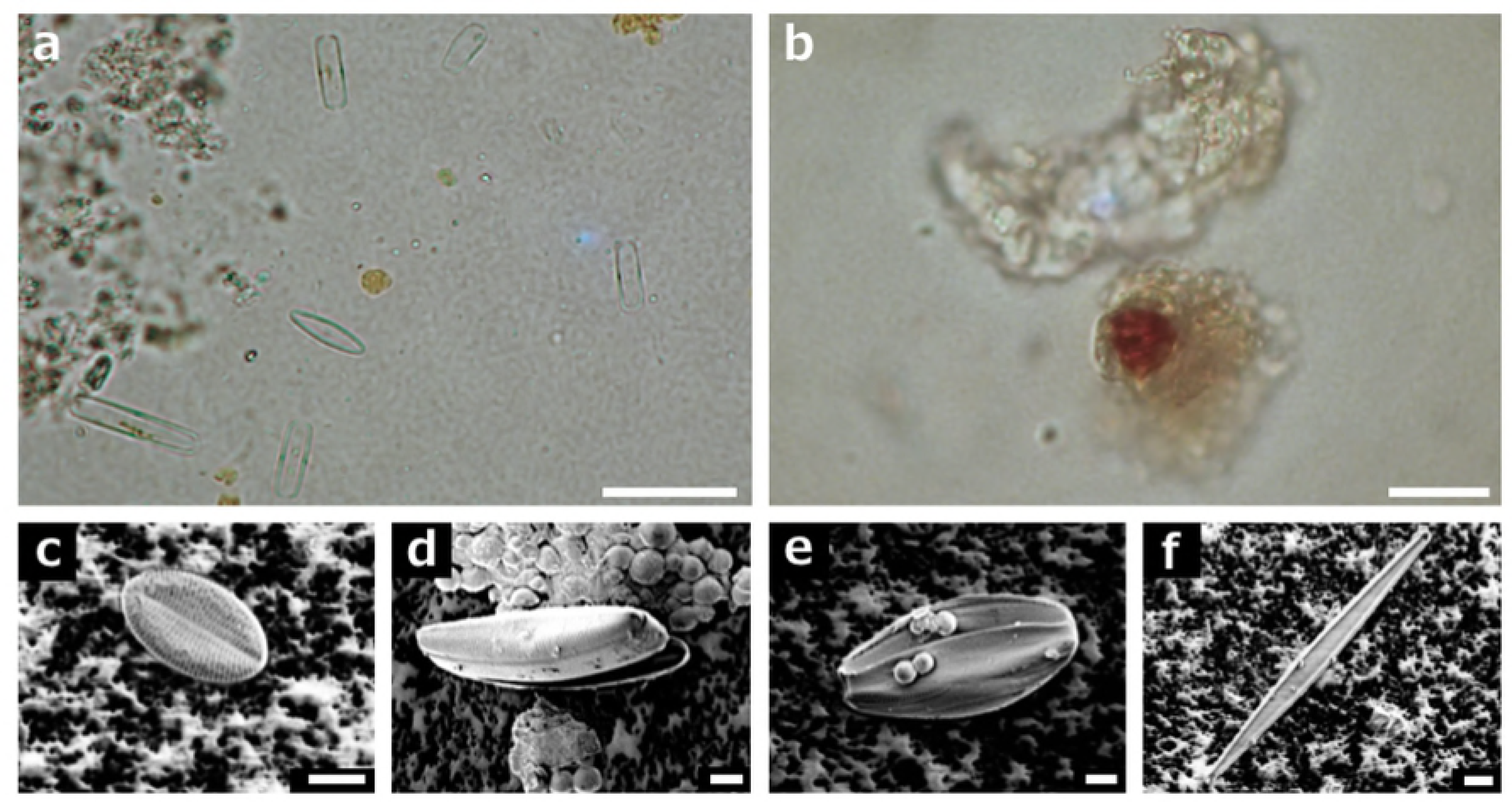
Stomach and intestinal contents of Fossarininae species observed using an optical (a–b) and an electron microscope (c–f). a, diatoms from the gastric contents of *R. eximia*; b, red algae from *R. eximia* observed using an optical microscope; c-f, Pennales diatoms. *Scale bar* **=** 100 μm (a–b), 5 μm (c–f).

**Table 3.** Organic matter seen in the gastric contents of each Fossarininae species.

|  | *F. picta* | *S. pulchella* | *B. iridescens* | *R. eximia* |
| --- | --- | --- | --- | --- |
| Pennales diatom | 158 | 19 | 49 | 135 |
| red alga | 5 | 2 | 3 | 2 |
| Nematoda | 0 | 0 | 0 | 1 |
| ossicle of sponge | 3 | 0 | 2 | 1 |
| Total | 163 | 21 | 52 | 138 |

## Discussion

An extensive survey of Fossarininae snails clarified that limpet-like *R. eximia* inhabit the lower intertidal zones of exposed rock reefs, but not protected reefs. Thus, we obtained sufficient information to differentiate the microhabitats of all four genera in Fossarininae. To determine why *Roya* snails inhabit only exposed rock reefs, we compared the macrobenthic and macrophytic communities of exposed and protected rock reefs at two sites. The results of this comparison suggest that both exposed and protected reefs were covered primarily by encrusting and branching red algae and branching brown algae, while algal coverage was significantly higher in exposed reefs than in protected reefs (Table 1, Fig 3). The low algal coverage of protected reefs may be caused by sedimentation on rock surfaces, as the growth of branching algae is hindered by sedimentation [24-25].

Another difference observed between exposed and protected reefs was in the faunal composition of sessile animals. At site E, exposed reefs were inhabited by large tetraclitid barnacles, while protected reefs were inhabited mainly by small balanid and chthamalid barnacles (Table 1). The faunal composition of barnacles is influenced by the availability of plankton, which is affected by wave action [26]. Although the density of barnacles differed significantly between sites A and E, tetraclitid barnacles always inhabited only the exposed rock reefs, and the vacant shells of these barnacles were utilized as refugia by *Roya* snails (Fig 5i).

The composition of mobile fauna was influenced by wave strength. Species richness and density of mobile organisms were generally higher for exposed reefs than protected reefs. *Montfortula picta* is a snail in the originally coil-less family Fissurellidae (Vetigastropoda), with a shell that bears a remarkable resemblance to that of *R. eximia*. *M. picta* was observed only on the exposed reef, and not on the protected reef, at both sites A and B, as was also observed for *R. eximia.* The coexistence of morphologically similar, phylogenetically distant snail species suggests that the limpet-like shell morphology is a product of convergence due to adaptation to life on exposed rock reefs.

*R. eximia* is unique in having a zygomorphic limpet-like shell despite belonging to the Trochidae, which generally have round, coiled shells. *R. eximia* belongs to subfamily Fossarininae, and all four recorded Fossarininae species were observed in this study. *S. pulchella*, which have somewhat flattened and loosely coiled shells, live in crevices. Therefore, it is possible that their flat shell is adaptive to narrow spaces. *B. iridescens*, which have flat limpet-like shells, live exclusively in the pits of sea urchins, and their flat shell is thought to be adaptive to life within the narrow free space of that habitat [22]. *Broderipia* snails have three pairs of lateral tentacles, which are longer than those of *Roya* snails. These long lateral tentacles may be useful for monitoring the position of their host sea urchin. On the other hand, the habitat of *R. eximia* is on rock surfaces that are constantly exposed to violent waves, and their shell morphology is advantageous for survival in wave-swept habitats for the following reasons. First, limpet-like shells are more tolerant of strong waves than coiled shells [27]. Second, limpet-like shells with large apertures promote the development of a highly contracted soft body with a large foot sole, which would confer strong clinging ability [4, 28]. Third, the radial ribs of the shell (Fig 7j–k) may reinforce structural integrity, defensive capability against carnivores or light interception. Limpets living in areas exposed to direct sunlight tend to have more ribs and nodules on the shell than those living in shaded areas, which serve to radiate heat [29], while shell ornamentation, including spines and ribs, offers protection against drilling carnivores [30-31].

*F. picta* possesses an operculum, which can be used for protection by shutting the shell aperture (Fig 7e). However, the operculum of *S. pulchella* is too small to cover the aperture, and *B. iridescens* and *R. eximia* have lost their opercula. Most gastropods with limpet-shaped shells or loosely coiled auriform shells have secondarily lost their opercula, and hard substrata substitute for the operculum [32]. Our results also suggest that loss of operculum has occurred during the evolutionary process of the loss of coiling in Fossarininae.

According to the observations of the diurnal behavior of *Roya*, *Roya* snails creep along rock surfaces to graze on periphytic diatoms and encrusting red algae at low tide, and then retreat into vacant shells of tetraclitid barnacles at high tide without migrating vertically through the intertidal zone. This behavioral pattern contrasts with many non-homing limpets that follow the wash zone of rocky intertidal zones. This vertical movement is considered to prevent attacks from aquatic predators, and simultaneously to minimize desiccation stress caused by exposure to air [33-34]. In contrast, *Roya* snails minimize predation pressure by moving over wave-swept rock surfaces at low tide and hiding in vacant shells of barnacles at high tide, rather than following the wash zone. Wave-swept rock surfaces are free from carnivorous muricid snails irrespective of tide level due to the violent waves. Although submerged rock surfaces are potentially vulnerable to carnivorous fishes at high tide, the vacant shells of barnacles act as refugia from predatory fishes.

Irrespective of shell morphology differences among the four Fossarininae species, their radulae were largely similar. In particular, *Synaptocochlea*, *Broderipia* and *Roya* have very similar radulae with sharp serrations, while *Fossarina* have spatula-like radula with shovel-like central teeth that have no serrations (Fig 9). Also, a comparison of gastric contents revealed that all Fossarininae species graze primarily on pennate diatoms on the surface of rocks, and the diatom species compositions were generally similar among the four Fossarininae species. These results suggest that the differing shell morphologies were not influenced by diet or foraging habits

Superimposing these observations on the phylogenetic tree of Fossarininae obtained by Williams (2010) [21], the evolutionary progression of shell morphology emerges (Fig 11). The basal lineage of Fossarininae, *Fossarina,* which inhabits open rock surfaces in protected reefs and crevices or vacant barnacle shells in exposed rock reefs, have typical coiled shells. In their sister clade (*Synaptocochlea + Broderipia + Roya*), found on exposed rock reefs, flattening of the shell likely occurred as an adaptation to life in refugia in wave-swept rock reefs.

**Fig 11.**
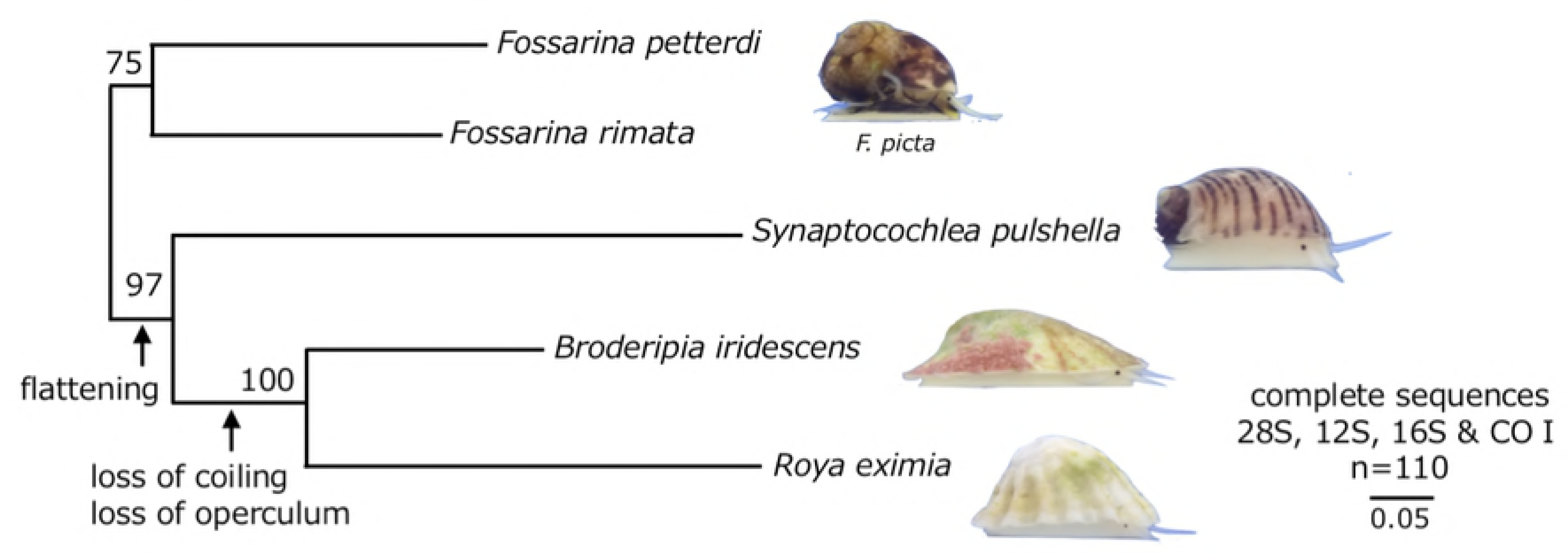
The progression of flattening and loss of coiling in subfamily Fossarininae overlaid with the phylogenetic tree obtained by Williams et al. (2010).

In the clade comprised of *Broderipia* and *Roya*, evolution of a completely zygomorphic limpet-shaped shell has occurred, although selective pressures may differ between the two genera. In *Broderipia*, the extremely flattened limpet-like shell is presumed to be an adaptation to the narrow free space offered by the pit of its host sea urchin. In *Roya*, the non-flat limpet-like shell with radial ribs is considered to be an adaptation to life moving between wave-swept rock surfaces and refugia of vacant barnacle shells. The coil-less limpet-like shell of *Roya* caused loss of coiling and contraction of the soft body, acquisition of a zygomorphic flat body, expansion of the foot sole, and loss of the operculum, all of which would have enhanced tolerance of strong waves and the power required to cling to rock surfaces. The factors that promoted the evolution of a limpet-like shell have been proposed to include the transition from mobile life to sessile life (e.g. Morton and Jones 2001) and advancing into a wave-exposed area (Branch 1985). Our results show that the evolution of limpet-like shells occurs under various selective pressures in coiled snail lineages. One intriguing topic for future research is determining the genes mutations that have contributed to the evolution of limpet-like shells in multiple lineages and various marine environments.

## Acknowledgements

We are grateful to all the staffs of the Seto Marine Biological Laboratory and Shirahama Aquarium for supporting our survey; and Y. Henmi and K. Kondo of Kyoto University for field assistance.

## Footnotes

The English in this document has been checked by at least two professional editors, both native speakers of English. For a certificate, please see: http://www.textcheck.com/certificate/Sk1Wmf

## References

[1] Eagar RMC. Shape and function of shell - Comparison of some living and fossil bivalve molluscs. Biological Reviews of the Cambridge Philosophical Society. 1978;53(2):169–210. doi: 10.1111/j.1469-185X.1978.tb01436.x. PubMed PMID: WOS:A1978FC64800001.

[2] Palmer AR. Fish predation and the evolution of gastropod shell sculpture – experimental and geological evidence. Evolution. 1979;33(2):697–713. doi: 10.2307/2407792. PubMed PMID: WOS:A1979HE57800016.

[3] Branch GM. Limpets: evolution and adaptation. The Mollusca. 1985;10:187–220.

[4] Branch GM. The biology of limpets: physical factors, energy flow, and ecological interactions. 1981.

[5] Lesicki A. Checklist of gastropod species referred to the order Cocculiniformia Haszprunar, 1987 (Gastropoda: Cocculinoidea et Lepetelloidea) with some remarks on their food preferences. Folia Malacologica. 1998;6(1-4):47–59.

[6] McLean JH. Cocculiniform limpets (Cocculinidae and Pyropeltidae) living on whale bone in the deep sea off California. Journal of Molluscan Studies. 1992;58:401–14. doi: 10.1093/mollus/58.4.401. PubMed PMID: WOS:A1992JZ46300006.

[7] Bates AE. Feeding strategy, morphological specialisation and presence of bacterial episymbionts in lepetodrilid gastropods from hydrothermal vents. Marine Ecology Progress Series. 2007;347:87–99. doi: 10.3354/meps07020. PubMed PMID: WOS:000250978500008.

[8] León-Cisneros K, Mazariegos-Villarreal A, Miranda-Saucedo CM, Argumedo-Hernández U, Siqueiros-Beltrones D, Serviere-Zaragoza E. Diet of the volcano keyhole limpet Fissurella volcano (Gastropoda: Fissurellidae) in subtropical rocky reefs of the Baja California Peninsula. Pacific Science. 2017;71(1):57–66.

[9] Herbert DG. Foraminiferivory in a puncturella (Gastropoda Fissurellidae). Journal of Molluscan Studies. 1991;57:127–40. doi: 10.1093/mollus/57.1.127. PubMed PMID: WOS:A1991EZ31400012.

[10] Kano Y, Chiba S, Kase T. Major adaptive radiation in neritopsine gastropods estimated from 28S rRNA sequences and fossil records. Proceedings of the Royal Society B-Biological Sciences. 2002;269(1508):2457–65. doi: 10.1098/rspb.2002.2178. PubMed PMID: WOS:000180108400011.

[11] Morton B, Jones DS. The biology of Hipponix australis (Gastropoda : Hipponicidae) on Nassarius pauperatus (Nassariidae) in Princess Royal Harbour, Western Australia. Journal of Molluscan Studies. 2001;67:247–55. doi: 10.1093/mollus/67.3.247. PubMed PMID: WOS:000171060200001.

[12] Collin R. The utility of morphological characters in gastropod phylogenetics: an example from the Calyptraeidae. Biological Journal of the Linnean Society. 2003;78(4):541–93. doi: 10.1046/j.0024-4066.2002.00166.x. PubMed PMID: WOS:000182544500010.

[13] Willan RC. A review of diets in the Notaspidea (Mollusca: Opisthobranchia). Journal of the Malacological Society of Australia. 1984;6(3-4):125–42.

[14] Walsby J, Morton J, Croxall J. The feeding mechanism and ecology of the New Zealand pulmonate limpet, Gadinalea nivea. Journal of Zoology. 1973;171(2):257–83.

[15] Bano A, Ayub Z, Siddiqui G. Fatty Acid Composition of Three Species of Siphonaria (Gastropoda: Pulmonata) in Pakistan. Pakistan Journal of Zoology. 2014;46(3):813–8. PubMed PMID: WOS:000338176400028.

[16] Blinn W, Truitt RE, Pickart A. Feeding ecology and radular morphology of the freshwater limpet Ferrissia fragilis. Journal of the North American Benthological Society. 1989;8(3):237–42.

[17] Fretter V, Graham A. British prosobranch molluscs. Their functional anatomy and ecology. British prosobranch molluscs Their functional anatomy and ecology. 1962.

[18] Schiaparelli S, Cattaneo-Vietti R, Chiantore M. Adaptive morphology of Capulus subcompressus Pelseneer, 1903 (Gastropoda: Capulidae) from Terra Nova Bay, Ross Sea (Antarctica). Polar Biology. 2000;23(1):11–6.

[19] Egloff DA, Smouse DT, Pembroke JE. Penetration of the radial hemal and perihemal systems of *Linxhia Laevigata* (Asteroidea) by the proboscis of *Thyca crystallina*, an ectoparasitic gastropod. Veliger. 1988;30(4):342–6. PubMed PMID: WOS:A1988Q265600002.

[20] Morton B. Partnerships in the sea: Hong Kong's marine symbioses: Kent State University Press; 1989.

[21] Williams ST, Donald KM, Spencer HG, Nakano T. Molecular systematics of the marine gastropod families Trochidae and Calliostomatidae (Mollusca: Superfamily Trochoidea). Molecular Phylogenetics and Evolution. 2010;54(3):783–809. doi: 10.1016/j.ympev.2009.11.008. PubMed PMID: WOS:000275176500010.

[22] Yamamori L, Kato M. The macrobenthic community in intertidal sea urchin pits and an obligate inquilinism of a limpet-shaped trochid gastropod in the pits. Marine Biology. 2017;164(3). doi: 10.1007/s00227-017-3091-3. PubMed PMID: WOS:000395180300024.

[23] National Institute of Advanced Industrial Science and Technology. 2011 [cited 2018 Apr 22]. Available from: https://gbank.gsj.jp/seamless/index_en.html

[24] Airoldi L. Roles of disturbance, sediment stress, and substratum retention on spatial dominance in algal turf. Ecology. 1998;79(8):2759–70. doi: 10.1890/0012-9658(1998)079[2759:rodssa]2.0.co;2. PubMed PMID: WOS:000077501000014.

[25] Airoldi L, Cinelli F. Effects of sedimentation on subtidal macroalgal assemblages: An experimental study from a Mediterranean rocky shore. Journal of Experimental Marine Biology and Ecology. 1997;215(2):269–88. doi: 10.1016/s0022-0981(96)02770-0. PubMed PMID: WOS:A1997XJ16400007.

[26] Akester RJ, Martel AL. Shell shape, dysodont tooth morphology, and hinge-ligament thickness in the bay mussel Mytilus trossulus correlate with wave exposure. Canadian Journal of Zoology-Revue Canadienne De Zoologie. 2000;78(2):240–53. doi: 10.1139/cjz-78-2-240. PubMed PMID: WOS:000085820200010.

[27] Denny M. Biology and the mechanics of the wave-swept environment: Princeton University Press; 2014.

[28] Vermeij GJ. A natural history of shells: Princeton University Press; 1995.

[29] Vermeij GJ. Morphological patterns in high-intertidal gastropods – adaptive strategies and their limitations. Marine Biology. 1973;20(4):319–46. doi: 10.1007/bf00354275. PubMed PMID: WOS:A1973Q477700006.

[30] Alexander RR, Dietl GP. The fossil record of shell-breaking predation on marine bivalves and gastropods. Predator—Prey Interactions in the Fossil Record: Springer; 2003. p. 141–76.

[31] Vermeij GJ. Biogeography and adaptation: patterns of marine life: Harvard University Press; 1978.

[32] Checa AG, Jimenez-Jimenez AP. Constructional morphology, origin, and evolution of the gastropod operculum. Paleobiology. 1998;24(1):109–32. PubMed PMID: WOS:000072055800008.

[33] Davies MS, Edwards M, Williams GA. Movement patterns of the limpet Cellana grata (Gould) observed over a continuous period through a changing tidal regime. Marine Biology. 2006;149(4):775–87. doi: 10.1007/s00227-006-0258-8. PubMed PMID: WOS:000238683900007.

[34] Iwasaki K. Factors affecting individual variation resting site fidelity in the Patellid limpet, Cellana toreuma (Reeve). Ecological Research. 1992;7(3):305–31. doi: 10.1007/bf02347099. PubMed PMID: WOS:A1992KM64100011.

